# Molecular and spatiotemporal characterization of cells in murine atherosclerotic plaques

**DOI:** 10.1101/2024.09.04.611323

**Authors:** Pengbo Hou, Zhanhong Liu, Jiankai Fang, Ziyi Wang, Shisong Liu, Shiqing Wang, Gerry Melino, Peishan Li, Yufang Shi, Changshun Shao

**Affiliations:** The Third Affiliated Hospital of Soochow University, Institutes for Translational Medicine, State Key Laboratory of Radiation Medicine and Protection, Suzhou Medical College of Soochow University, 215123 Suzhou, China; Department of Experimental Medicine, TOR, University of Rome Tor Vergata, 00133 Rome, Italy

**Keywords:** Atherosclerosis, Imaging mass cytometry, Spatial landscape, scRNA-seq, Plaque cells, Inflammation

## Abstract

**Objective:** Single-cell technologies have revolutionized our understanding of the phenotypic and transcriptional diversity of aortic leukocytes in atherosclerotic humans and mice. However, enzymatically dissociated tissues lose the spatial context of plaque cells *in situ*. Here we utilized imaging mass cytometry (IMC) combining with single-cell RNA sequencing (scRNA-seq) to characterize the spatial distribution dynamics, phenotypic transitions, metabolic and functional phenotypes, and the intercellular interaction networks of plaque cells during atherosclerotic progression. Additionally, the dynamic immune landscape of circulating leukocytes associated with atherosclerosis was characterized using cytometry of time of flight (CyTOF).

**Approach and Results:** A highly multiplexed IMC panel with 33 metal-conjugated antibodies was designed to generate 11 highly multiplexed histology images of aortic root tissues from *ApoE*^-/-^ mice on high-fat diet at different stage of atherosclerosis. Using histoCAT, we identified 8 principal cell subtypes with distinct phenotypic and geographic dynamics. Furthermore, IMC-defined cell subsets partially corresponded to scRNA-seq-annotated aortic cell subtypes, including 4 macrophage subsets, neutrophils, smooth muscle cells (SMCs) and SMC-derived SEMs (Stem cell, endothelial cell and macrophage-like cell). Activation of inflammatory pathways, increased oxidative phosphorylation and augmented osteoclast differentiation were observed in macrophage populations, SMCs and SEMs from an early stage to advanced stage of atherosclerosis. Notably, cell neighborhood analysis by IMC uncovered multifaceted cell-cell interactions within the plaque, in particular in neutrophil-mediated interactions with smooth muscle cells and macrophages, which were confirmed by ligand-receptor interactions based on scRNA-seq data.

Additionally, characterization of the peripheral immune cells by CyTOF revealed an increased ratio of myeloid cells to lymphocytes, and certain neutrophil and monocyte subpopulations also exhibited enhanced lipid metabolism and glycolysis as well as activated inflammatory signaling.

**Conclusion:** This study provides a dynamic spatiotemporal landscape of atherosclerotic lesions and peripheral leukocytes. The new information based on IMC may help understand atherosclerotic pathology and develop novel therapeutic strategies.

## 1. Introduction

Atherosclerosis, a lipid-driven chronic inflammatory disease, is characterized by leukocyte infiltration and lipid-core formation within the intima of large arteries. Atherosclerotic cardiovascular diseases (CVD), such as myocardial infarction and stroke, have emerged as one of leading causes in human mortality globally^1^. The RESCUE clinical trial by interleukin (IL)-6 inhibition^2^, and the CANTOS clinical trial by blocking IL-1β^3^ support that anti-inflammatory interventions are effective in reducing cardiovascular events. Accordingly, deciphering the landscape of immune cells and their niche during atherosclerotic progression will enable us to distinguish between pathogenic and reparative subpopulations, thus facilitating the development of targeted therapeutic strategies.

Applications of advanced single-cell technologies, such as single-cell RNA sequencing (scRNA-seq) and mass cytometry by time-of-flight (CyTOF), have helped uncover the heterogeneity of aortic immune cells and their distinct biological features in atherosclerotic humans and mice. Aortic macrophages can be classified into distinct subtypes by scRNA-seq analysis, including resident-like macrophages, TREM2^hi^ (foamy) macrophages and inflammatory macrophages^4-6^. Additionally, the heterogeneity of non-immune cells (including smooth muscle cells, endothelial cells and fibroblasts) and their phenotypic switch in atherosclerosis have been revealed in different studies^7-9^. However, these two technologies rely on tissue dissociation, and thus do not provide spatial information of different types of cells in the plaque, such as geolocation and neighboring cell relationships. In addition, existing bioinformatic tools used for analyzing intracellular communication primarily relying on scRNA-seq data, such as CellPhoneDB^10^, NicheNet^11^, and CellChat^12^, are dependent on ligand-receptor gene expression information rather than cell location. Indeed, such modeling does not accurately reflect the in vivo conditions, given that cells with complementary ligands and receptors may be physically distant from each other within the tissue microenvironment.

The emergence of imaging mass cytometry (IMC) using metal-conjugated antibody has addressed the aforementioned limitations and enabled us to obtain comprehensive information from proteins to cell to environmental context by highly multiplexed epitope-based tissue imaging^13^. IMC has been widely used for deep studies of tissue and tumor biology at the single-cell level with spatial resolution in tissues^14-16^, but a comprehensive characterization of atherosclerotic plaque at different stages of disease using IMC has not been conducted.

In this study, we employed IMC to simultaneously examine the expression of 33 proteins within murine atherosclerotic plaques at early, intermediate and advanced stages. We interrogated the dynamic spatial landscape of atherosclerotic plaques with distinct topography of immune cells, stromal cells and smooth muscle cells, which were substantiated by parallel aortic cell subtypes identified by scRNA-seq. Topological analysis of plaque environment uncovered a crucial role of neutrophils in atherogenesis by interacting with smooth muscle cells. Moreover, the identified interaction pairs based on cell localization were also confirmed by the mode dependent on ligand-receptor interactions. By leveraging the high-throughput advantages of scRNA-seq, we unveiled highly enriched pathways of oxidative phosphorylation, osteoclast differentiation, and inflammation-related pathways in macrophages and smooth muscle cells along with the disease progression. Notably, CyTOF with 32 metal-labeled antibodies revealed increased abundance of inflammatory cells in peripheral blood that is associated with atherosclerotic lesions, further confirming that atherosclerosis is a systematic inflammatory disease. Collectively, our study revealed a comprehensive and longitudinal landscape of atherosclerotic plaques in mouse model.

## 2. Materials and Methods

Expanded methods are available in the Supplementary material

### 2.1 Animal models

Apolipoprotein E deficient (*Apoe^-/-^*) male mice of C57BL/6J background (6-8 weeks) were obtained from Gempharmatech Co., Ltd (Nanjing, China), and were maintained in a specific pathogen-free facility of the Laboratory Animal Center of Soochow University. Mice were randomly separated into four groups (n = 5 per groups). After a two-week adaptation period, the mice were provided with either chow diets or high-fat diets containing 40 kcal % of fat (Product ^#^12108C, SHUYISHUER BIO, Changzhou, China) at varying durations (8, 11, 19, 26 weeks). Animal care and experiments were approved by and in full compliance with the guide for the care and use of the Institutional Animal Care and Use Committee of Soochow University (SUDA20210916A02).

### 2.2 Sample collection and processing

#### Blood

After mice were anesthetized with isoflurane, blood samples (about 800 μL per mouse) were collected from the orbital vein followed by cervical dislocation. The collected blood was then processed to remove red blood cells using a red blood cell lysis buffer (Solarbio, China). The obtained leukocytes were washed with pre-cooled PBS followed by metal-labelled antibodies staining.

#### Aorta

After cervical dislocation, the mouse vasculature was perfused by cardiac puncture with chilled PBS to remove blood from all vessels. Subsequently, the aorta was carefully harvested using a stereomicroscope after pulling off all surrounding adipose tissue. Surgically excised aortas were collected in tissue storage solution (130-100-008, Miltenyi Biotec). Aortas were digested on the basis of previous description^4^ with a minor modification.

#### Aortic root

After removal of the apex portion of the heart, the remaining part containing aorta root was embedded in OCT Tissue TEK and rapidly frozen in liquid nitrogen, then stored at -80 °C in a freezer before tissue cutting.

#### Aortic CD45^+^ cell isolation for scRNA-seq

To sort leukocytes from single-cell suspension of aortas, we utilized a CD45 MicroBead-dependent approach (130-052-301, Miltenyi Biotec) to minimize sorting time and maintain high cell viability. The detailed protocol was provided by Miltenyi Biotec. For more details see supplemental materials.

### 2.3 The design of panels for IMC and CyTOF

Using the panel design helper (Version: 2.83) provided by Fluidigm, we designed two panels for IMC (Table 1) and CyTOF (Table 2). Table 1 and 2 were displayed in supplemental materials.

### 2.4 Mass cytometry antibody conjugation

For IMC and CyTOF experiments, directly metal-conjugated antibodies were purchased from Fluidigm. Alternatively, purified antibodies were labelled with the Maxpar X8 Multi-metal labeling Kit (Fluidigm) according to the manufacturer’s instructions. Conjugation efficiency was verified by NanoDrop One (ThermoFisher). All antibody concentrations were titrated.

### 2.5 Mass cytometry

#### Viability staining for blood leukocytes

Cells were washed in PBS (without Ca^2+^ and Mg^2+^) and stained Cisplatin (Fluidigm) in a final concentration of 5 mM. Cells were washed and stained with the antibody cocktail listed in Table 2. Antibodies were prepared in cell staining buffer (Fluidigm). After staining, cells were washed and fixed with 1.6% Formaldehyde (FA) buffer (prepared by 16% FA, Thermo fisher scientific) to improve the detection of intracellular protein. For cell identification, cells were washed in staining buffer and stained with premixed cell intercalation (prepared to a final concentration of 125 nM Iridium in 1 × Fix and Permeabilization buffer, Fluidigm). Cells were washed in staining buffer and ultra-pure water (Fluidigm) to remove buffer salts. Cells were next resuspended in cell acquisition solution (CAS) buffer with a 1:10 dilution of EQ Four Element Calibration beads (Fluidigm) and filtered through a 40 μm nylon mesh filter cap. Samples were acquired on a Helios CyTOF Mass Cytometry (Fluidigm) at an event rate of 500 events/second.

#### IMC staining for aortic root sections

Frozen aortic root slides underwent immunostaining following the manufacturer’s instructions. In brief, the slides were washed with 1× PBS and incubated in PBS containing 3% bovine serum albumin (BSA) for 45 minutes. Subsequently, the slides were stained with a cocktail of metal-labeled antibodies at optimized dilutions overnight at 4 °C. Afterward, the slides were washed with 0.2% Triton X-100 and 1× PBS. Prior to IMC acquisition, Cell-ID Intercalator-Ir (Fluidigm) at a dilution of 1:400 was utilized to counterstain slides in 1× PBS for 30 minutes at room temperature. The slides were then rinsed for 5 minutes with distilled water and air-dried.

### 2.6 CyTOF and IMC data analysis

#### For CyTOF data analysis

The raw file of CyTOF data were generated in .fcs file format by Helios software. These .fcs files were then uploaded to Cytobank (https://premium.cytobank.org/cytobank) for all gating that removes EQ Four Element Calibration beads and cohesive and dead cells. The gated .fcs files were transferred from Cytobank to R (Version 4.2.2) and were subjected to an automated dimensionality reduction algorithm *t*SNE and Phenograph through Bioconductor package Cytofkit (Version 3.6).

#### For IMC data analysis

An imaging mass cytometer (Fluidigm, Hyperion) was used to scan the prepared aortic sections to generate multiplexed images. To segment image data into single-cell data, we used CellProfiler software (Version 4.2.4) to obtain mask files, which were fed into histoCAT software (Version 1.73) for *t*SNE and PhenoGraph analyses and cell neighborhood analysis^17^ (https://bodenmillergroup.github.io/histocat-web/).

### 2.7. scRNA-seq and analysis

scRNA-seq of sorted aortic leukocytes from ApoE^-/-^ mice was performed using 10 × Genomics platform. The library synthesis and RNA-sequencing, and bioinformatic analysis were completed by the Gene Denovo (Guangzhou, China). For detailed methodology see supplemental materials.

### 2.8 Statistics

Raw data obtained from FACS and histoCAT were copied into GraphPad software (Version 9.5.1, La Jolla, USA). One-way ANOVA analysis of Dunnett’s multiple comparisons test or Tukey’s multiple comparisons test was used to identify differences among three or more groups. The data were represented as the means ± SEMs. A *p* value <0.05 indicated statistical significance.

## 3. Results

### 3.1 Establishment of IMC analysis for atherosclerotic plaques

To map the dynamic landscape of plaque microenvironments during atherosclerosis progression, *ApoE^-/-^* mice were subjected to chow diets or high-fat diets (HFD) for different intervals (8, 19 and 26 weeks) to simulate different stages of atherosclerosis (Figure 1A). The aortic root sections were stained by a highly multiplexed metal-conjugated antibody panel of 33 markers, including markers for immune cells, smooth muscle cells (SMCs) and stromal cells, markers for lipid metabolism (CPT1A and CD36), glycolysis (GLUT1 and HK2), proliferation (Ki-67) and apoptosis (cleaved caspase-3) (Supplement materials, Table 1). We also performed H&E staining of aorta roots to evaluate the histological features at different plaque stages. We found increased areas of neointima and necrosis along with atherosclerotic progression, and detected hemorrhage in advanced plaque (Figure 1B). Based on H&E staining, a total 11 panoramic images of aorta root sections derived from 4 groups, Chow diet (n = 2), HFD-8W (n = 3), HFD-19W (n = 3) and HFD-26W (n = 3), were selected for laser scanning by Hyperion-imaging system, and the high-dimensional histopathology images were subsequently analyzed by IMC pipeline. The major cell subpopulations within the plaque were classified based on their expression of specific cell markers and histopathology localization (Figure 1C-D), including macrophages (CD68^+^), monocytes (Ly6C^+^, CD68^-^, Ly6G^-^), fibroblasts (PDGFRα^+^, Vimentin^+^, CD90.2^+^), SMCs (α-smooth muscle actin, αSMA^+^) and neutrophils (Ly6G^+^, MPO^+^) (Figure 1E). Rare T cells (CD3^+^, CD90.2^+^) were only detected in the adventitial regions of blood vessels (Figure 1E panel4).

**Figure 1.**
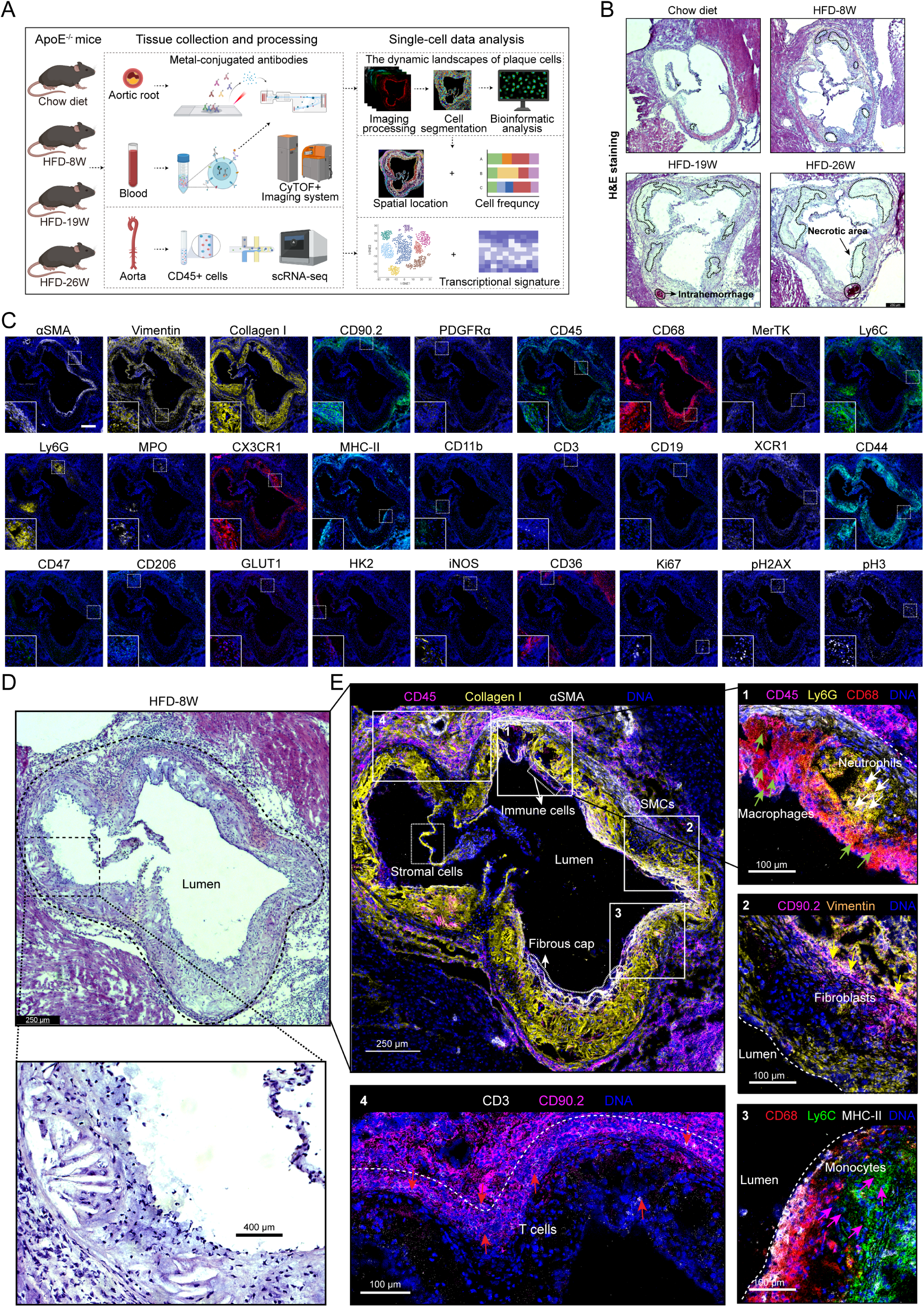
Establishment of workflow for mapping the dynamic landscape of plaque cells. **(A)** This workflow entails the utilization of IMC analysis, scRNA-seq analysis, and mass cytometry analysis at the single-cell level for aortic root sections, aortas, and circulating immune cells obtained from ApoE-KO mice at various disease stages. Aorta root sections and blood samples were subjected to multiplex staining, and data were acquired using cytometry by time-of-flight (CyTOF) technology. IMC pipeline includes cell segmentation, cell dimensionality reduction, identification of cell subsets and spatial geolocation. **(B)** Evaluation of histological features of plaque based on H&E images. The dashed line represents the necrotic regions, while the solid line represents intraplaque hemorrhage. Scale bar, 250 μm. **(C)** Representative mass cytometry images of aortic root from HFD-8W group. Scale bar, 250 μm. **(D)** Representative H&E staining image of aortic root from HFD-8W group. **(E)** Identification of cell types based on the expression of cell markers. CD45 (purple), Collagen I (yellow), αSMA (white), DNA (bule) were used to specify immune cells, stromal cells, SMCs and fibrous cap. **1)** CD68 (red), Ly6G (yellow), were used to specify neutrophils and macrophages. **2)** CD90.2 (purple), Vimentin (yellow), were used to identify fibroblast. **3)** CD68 (red), Ly6C (green), and MHC-II (white) were used to identify monocytes. **4)** CD3 (white), CD90.2(purple), were used to identify T cells. Scale bar, 100 μm.

### 3.2 Dynamic landscape of plaque cells

Next, we attempted to investigate the spatial distribution of plaque cells and their dynamic changes during atherosclerotic progression. The *t*SNE algorithm and PhenoGraph analysis were applied to the segmented 89,381 cells with marker expression information through histoCAT^17^. We identified 24 distinct cell clusters, which were subsequently visualized on a *t*SNE plot (Figure 2A) and observed significant landscape changes in the cellular composition of plaques at various disease stages (Figure 2B). These clusters were further annotated based on the variation in marker expression intensity among different cell subpopulations (Figure 2C-E, supplemental Fig S1A) and their respective spatial locations. The main features of the 24 cell subsets within the plaque microenvironment are as follows:

**Figure 2.**
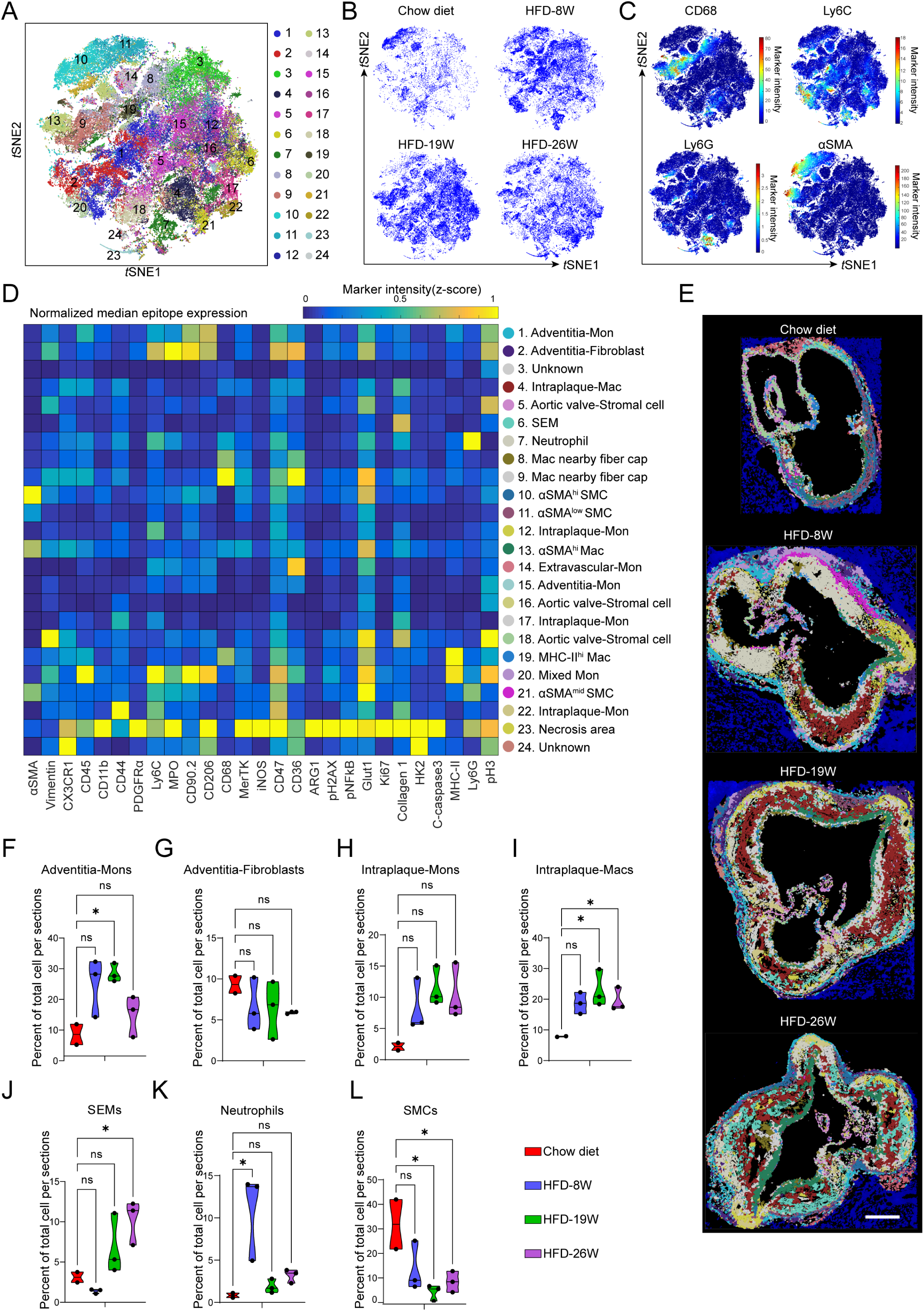
IMC reveals the spatiotemporal changes of plaque cells. **(A)** Combined *t*SNE plot of 24 cell clusters are displayed for representative all plaque cells from the scanned 11 images. **(B)** *t*SNE plots of Chow diet, HFD-8W, HFD-19W and HFD-26W, respectively. **(C)** *t*SNE plots of cell markers. **(D)** Heatmap showing average expression of all markers across cell types in the atherosclerotic aortic root using the panel of isotope-conjugated antibodies. **(E)** The spatial landscape of plaque cells at various stage of atherosclerosis are shown, scale bar 250 μm. **(F-L)** Violin plot showing the percentage of the same cell subtype (pooled) within all cell subtypes in the entire sections at different stages of disease. Data are presented as mean ± SEM. Dots represent individual samples, n = 2-3, \**p*<0.05, \*\**p*<0.01, by one-way ANOVA with Dunnett’s multiple comparison test.

Clusters 1 and 15 were designated as Adventitia-Mon due to their localization in the adventitia of the aorta (Figure 2D and E). Cluster 1 also highly express CD206, reminiscent of the atheroprotective resident-like macrophages (high expression of *Lyve1*, *Cd209f*, *Cd206*, and low expression of *Il6*, *Tnf* and *Nfĸbia*) reported in previous studies^18,19^. Our scRNA-seq results suggest that the two clusters belong to the same cell type. Moreover, the low expression of Ki67 in Adventitia-Mon (cluster 1) indicates their low proliferation capacity. The two Adventitia-Mon subsets appeared to be more highly represented in the HFD-fed groups than in chow diet group, particularly in the HFD-19W group (Figure 2F, Supplemental Fig S1B).

Cluster 2, referred to as Adventitial-Fibroblast, showed no significant change in percentage of cell numbers (Figure 2G). These cells were characterized by positive expression of fibroblast markers (Vimentin and CD90.2) and high expression of myeloperoxidase (MPO), CD36, and phosphorylated histone H3 (Figure 2D).

Cluster 4, 8, 9, and 19, characterized by the positive expression of CD68 and CD45, were commonly categorized as Intraplaque-Mac subpopulations that were absent in Chow diet but were prominently present in the plaque of the HFD-19W and 26W groups (Figure 2I). Among these, cluster 4 exhibited low expression of CD68, MerTK, CD36, Glut1, and MHC-II, but had intermediate expression of Collagen Ⅰ. Cluster 8 showed moderate expression of CD68 and CD36 as well as low expression of Glut1. Cluster 9 possessed high expression of CD68, Glut1, and CD36, with moderate expression of MerTK and low expression of Ki67. Cluster 19 displayed high expression of MHC-II and Glut1, regarded as inflammatory macrophages (Figure 2D). Interestingly, the spatial proximity of cluster 8 and 9 to lipid-rich regions suggested that they may represent foamy macrophages (or TREM2^hi^ Mac) identified by scRNA-seq (Figure 2E).

Clusters 12, 17 and 22 that express Ly6C were designated as Intraplaque-Mon. The percentage of these clusters within each section exhibited an increasing trend during atherosclerotic progression (Figure 2H).

Cluster 7 was defined as neutrophils based on their co-expression of MPO and Ly6G. We further observed that the number of neutrophils reached its peak in the plaques of the HFD-8W mice (Figure 2E, K, and supplemental Fig S2B), which indicated a significant infiltration of neutrophils during the early stage of atherosclerosis. However, there was a reduction in the number of neutrophils in advanced plaques (Figure 2E and supplemental Fig S2B). These findings suggest that neutrophils may play a critical role in the initiation and early development of atherosclerotic plaques.

Cluster 6 was predominantly localized in the intima near the media region, and was increasingly represented in late-stage plaques (Figure 2E, J and supplemental Fig S2A). Based on literature^7^, plaque cells that highly express Collagen Ⅰ primarily include fibroblasts, SMCs, and SMC-derived SEMs. Since Cluster 6 cells highly expressed Collagen Ⅰ rather Vimentin or αSMA, they were classified as SEM.

Clusters 10, 11 and 21 were identified as SMCs based on their expression of αSMA and their predominant localization in the vascular media (Figure 2D and E). Furthermore, they became significantly less represented as atherosclerosis progressed (Figure 2L). Cluster 10 highly expressed αSMA and Glut1, and was more abundant in plaques from the Chow diet group compared to the HFD-fed group (Supplemental Fig S2C). Cluster 11, representing αSMA^low^ Glut1^mid^ SMC, was present in all stage of plaques, with a small subset of cluster 11 cells located near the lumen in the HFD-26W group (Supplemental Fig S2D). Cluster 21, which moderately expressed αSMA but highly expressed Glut1, was detected predominantly in the HFD-8W group (Supplemental Fig S2E). These data demonstrated the heterogeneity of SMCs in the plaques and their complicated dynamics during disease progression.

Cluster 13, located in the fibrous cap, was defined as αSMA^hi^-Mac based on its co-expression of αSMA and CD68, while their moderate expression of CX3CR1 and CD45 indicated their origin from hematopoietic stem cells rather than smooth muscle cells. Additionally, the high expression of GLUT1 and low expression of Ki67 in these cells suggest their metabolic preference of glycolysis and low proliferative activity. Moreover, they expressed Collagen Ⅰ and accounted for the majority of the cell population in the fibrous cap, possibly contributing to plaque stability. Notably, they basically remained unchanged in their locations during disease progression, and likely resided before the formation of plaque (Figure 2D and Supplemental Fig S2F).

### 3.3 Integrated functional analysis of plaque cell subpopulations by scRNA-seq

Because imaging mass spectrometry only allows the staining with a maximum of 40 metal-conjugated antibodies, it is challenging to accurately assess the functional and metabolic states of cell subsets within the plaque. Therefore, we employed scRNA-seq to better elucidate cell states of different plaque cell types during disease progression.

We utilized magnetic beads conjugated with the CD45 antibody to isolate all the viable CD45^+^ leukocytes from the aortas of male *Apoe*^−/−^ mice fed on a high-fat diet (HFD) for different time intervals (11, 19 and 26 weeks), respectively representing the intermediate (11W) and advanced stage (19, 26 W) of atherosclerotic lesion. We captured 13,813 cells from the HFD-11W, 8,303 cells from the HFD-19W, and 12,975 cells from the HFD-26W group. Gene expression data from all captured cells were extracted and subjected to bioinformatic analysis. Using unsupervised Seurat-based clustering (see methods section), the captured cells were classified into 22 distinct cell clusters as shown in UMAP (Uniform Manifold Approximation and Projection) plot (Supplemental Fig S3A). They were annotated based on their distinct expression profiles in combination with published single-cell transcriptomes studies for atherosclerotic lesions^4,7,18^ and established frameworks for defining myeloid cells (*Adgre1* encoding F4/80 antigen, *Itgax* encoding Cd11c, *Fscn1*, and *Fcgr1* encoding Cd64), lymphocyte lineages (Cd3e, Nkg7), and smooth muscle cells (*Acta* encoding *αSMA*, and *Myh11*), (Supplemental Fig S3B and C).

Macrophages, the largest cell population in the atherosclerotic aorta, include 5 subpopulations (Gpnmb^hi^-Mac, Inflam-Mac, Adven-Mac, Prolifer-Mac, and Mac2). Gpnmb^hi^-Mac (cluster 1, *Atp6v0d2*, *Gpnmb*, *Mmp12* was annotated as foamy macrophage or TREM2^hi^ macrophages^18^ (Supplemental Fig S3D). Inflam-Mac (cluster 2) (*H2-Eb1*, *Cd74* and *Clec4b1*) is likely involved in osteoclast differentiation, Th17 cell differentiation, and antigen processing and presentation according to the KEGG analysis (Supplemental Fig S3E). Advent-Mac (cluster 3) (*Cd209f*, *Lyve1*, *Cd209d*, *Fcna*, *Cd163*, and *Cbr2*) was defined as a resident-like macrophage subset in several published studies^18,19^. These macrophages are localized in the vascular adventitia and usually conduct endocytosis and lysosomal degradation^20^ (Supplemental Fig S3C and F). Prolifer-Mac (cluster 6: highly expressing *Ube2c*, *Birc5* and *Pclaf*) is implicated in proliferative process by gene ontology analysis.

Non-macrophage cell populations included neutrophils (cluster 4, represented by *S100a8*, *S100a9* and *Cxcr2*), monocytes (cluster 8, represented by *Chil3*, *Vcan* and *Ly6c2*, osteoclast differentiation and C-type lectin receptor signaling pathway) (Supplemental Fig S3C and H), B cells (cluster 16, represented by *Iglc1*, *Fcmr*, and *Ms4a1*), three clusters of dendritic cells (DC1, cluster 14 - *Gm43914*, *Xcr1* and *Clec9a*; DC2, cluster 7 - *Cd209a*, *Cd7* and *Klrd1* (Supplemental Fig S3C and G); DC3, cluster 15 – *Fcn1*, *Cacnb3* and *Ccr7*), CD8^+^ T cells (cluster 9 - *Xcl1*, *Nkg7* and *Gimap7*) (Supplemental Fig S3C and I), CXCR6^+^ T cells (cluster 12 -*Cxcr6*, *Trdc*, *Ly6g5b*, *Trgc1* and *Cd3g*), natural killer (NK) cells (cluster 17 -*Gata3*, *Klrg1*, *Calca* and *Tnfrsf18*), Mast cells (cluster 19 - *Cma11*, *Mrgprb1* and *Mcpt4*).

Due to the technical limitations with magnetic bead sorting, CD45 positive cells were not 100% pure and contained a small fraction of non-immune cells. The non-immune cells were subsequently identified as SMC-derived SEM (cluster 5- *Chl1*, *C7*, *Fbln* and *Serping1*, characterized by an intermediate cell state with multipotent differentiation potential towards macrophage-like cells, mesenchymal stem cells, and fibrochondrocytes^7^), SMCs (cluster 10 - *Itga8*, *Myh11* and *Tagln*), modulated SMCs (cluster 11 - *Ltbp2*, *Col8a1* and *Col6a3*), Osteochondrocytes (cluster 20 - *Acan*, *Ibsp*, and *Col11a1*), endothelial cells (EC, cluster 18 - *Esam*, *Ecscr* and *Ptprb*) (Supplemental Fig S3C).

To investigate the dynamic changes in cellular states during disease progression, we utilized a predefined gene set (Supplemental Table3) encompassing metabolic or functional states of plaque cells, including chemotaxis, cholesterol efflux, electron transport chain (ETC), glycolysis, oxidative phosphorylation (OXPHOS), apoptosis, necroptosis, NADPH oxidase, reactive oxygen species (ROS) production, phagocytosis, and the tricarboxylic acid cycle (TCA). Surprisingly, our findings revealed an overall upregulation of chemotaxis in plaque cells, indicating heightened cell migration responses (Figure 3A). Conversely, there was a notable reduction in cholesterol efflux and phagocytosis (Figure 3A). The dysregulated lipid metabolism and impaired efferocytosis may contribute to the persistence of excessive lipid accumulation and trigger sustained hyperinflammation, thus promoting the development of atherosclerosis. Additionally, we evaluated the distribution of cellular states in distinct cell clusters (Figure 3B). We found that Gpnmb^hi^ Macs (cluster 1) exhibited more active cellular characteristics such as glycolysis, OXPHOS, ETC, and phagocytosis, compared to Inflam-Macs and Adven-Macs (Fig 3B-F). Plaque neutrophils (cluster 4) exhibited a preference for NADPH oxidase and chemotaxis (Fig 3G-H).

**Figure 3.**
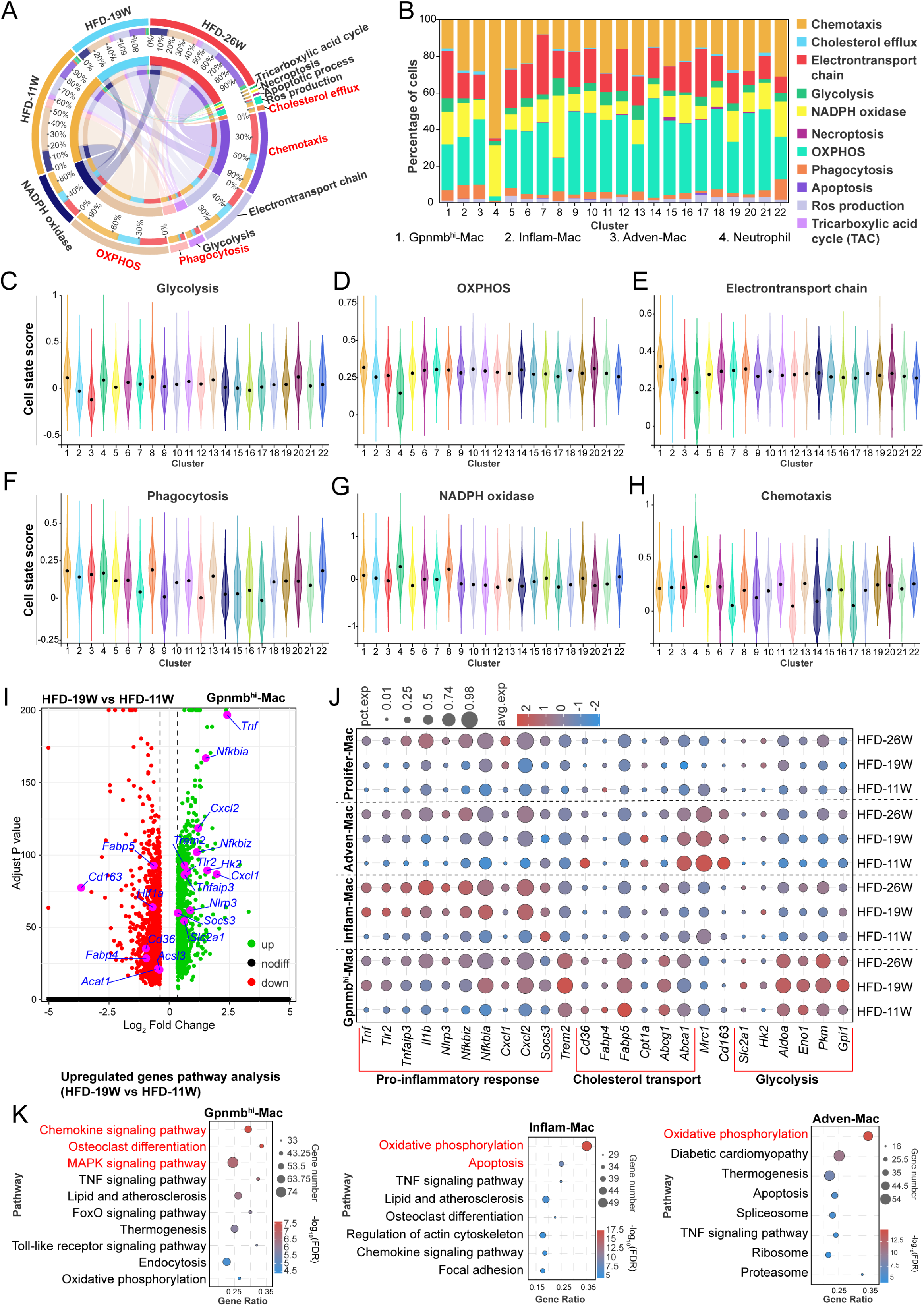
scRNA-seq characterization of cellular states and functional and metabolic phenotypes in macrophage populations during atherosclerosis. **(A)** Circus plot showing the overall changes of cell state of plaque cells at different stage of disease. **(B)** Stack diagram showing the distribution of different cell states in each cell cluster. **(C-H)** Violin plot representing the score of specific cell state in each cell cluster. Normalized data. **(I)** Volcano plot of representative upregulated or downregulated genes in Gpnmb^hi^-Mac (HFD-19W vs HFD-11W). Genes with adjusted *P* values <0.05 (red) are arranged by their *P* values and fold changes (log2). **(J)** Bubble plot of the expression of representative pro-inflammatory and metabolic genes in four macrophage subpopulations at different stage of disease. The size of each bubble represents the proportion of cells expressing that gene within the population. Larger bubbles indicate a higher proportion of cells expressing the gene. The color of the bubble, with a redder shade, represents a higher average expression of the gene within the cell subtype. **(K)** Upregulated genes (HFD-19 vs HFD-11W) enriched signaling pathways in three distinct macrophage subsets by KEGG analysis.

### 3.4 Functional and metabolic changes of macrophage subpopulations during atherosclerotic progression

As the predominant cell populations within the plaque, macrophages play a crucial role in plaque progression and stability. We therefore conducted a comprehensive analysis of the functional markers in distinct macrophage subpopulations. Our findings revealed significant upregulation of pro-inflammatory genes (*Tnf*, *Nfkbia*, *Nfkbiz* and *Nlrp3*) and notable downregulation of cholesterol transport genes (*Cd36*, *Fabp5* and *Abcg1*) and anti-inflammatory genes (*Mrc1* and *CD163*), in the Gpnmb^hi^- Mac subpopulation of HFD-19W group compared to HFD-11W group (Figure 3I). These phenotypes were similarly observed in other subpopulations of macrophages such as Inflam-Mac, Adven-Mac and Prolifer-Mac. Additionally, Gpnmb^hi^-Macs exhibited a significant enrichment of glycolysis-related genes (*Aldoa*, *Eno1*, *Pkm* and *Slc2a1*) in comparison to other macrophage populations, with their expression levels increasing during the disease progress, indicating the coexistence of hyperactive OXPHOS and glycolysis in the Gpnmb^hi^-Mac subpopulations during atherosclerosis (Figure 3J).

To further apprehend the dynamic functional changes in these macrophage subsets, we performed Kyoto encyclopedia of genes and genomes (KEGG) analyses on the upregulated genes of Gpnmb^hi^-Mac, Inflam-Mac, Adven-Mac and Prolifer-Mac (HFD-19 vs HFD-11W). As expected, upregulated genes in these macrophage subsets were similarly enriched in signaling pathways, including chemokine signaling pathway, tumor necrosis factor (TNF) signaling pathway, osteoclast differentiation and apoptosis, indicating that these 4 macrophage subsets possessed sustaining and anabatic inflammatory response and cell death process along with the disease progression (Figure 3K). While oxidative phosphorylation (OXPHOS) is generally considered to be preferred in anti-inflammatory macrophages, it was also enriched in plaque macrophage subgroups, especially in Inflam-Mac and Adven-Mac subpopulations. It may represent a feedback response to hyperinflammation and excessive lipid contents.

### 3.5 Functional and metabolic changes in plaque monocytes and neutrophils during atherosclerosis

Our IMC data have demonstrated that monocytes and neutrophils are implicated in the development of atherosclerotic plaques. We next ascertained whether the pro-inflammatory characteristics of monocytes and neutrophils were intensified as the disease advances. Through KEGG analysis, we observed that compared to the HFD-11W, upregulated genes in monocytes (HFD-19W) were enriched in apoptotic pathways and oxidative phosphorylation pathways (Figure 4A and B), and inflammatory genes such as *Il1β, Tnf, NFkbia* and *Nfkbiz* in monocytes were further increased as disease progressed (Figure 4C). Although previous *in vitro* experiments confirmed that enhanced mitochondrial OXPHOS in monocytes implies their differentiation into anti-inflammatory M2-like macrophages^21^, this theory has not been verified in monocytes or macrophages of atherosclerotic plaques, due to the intricate plaque microenvironment. Here, we propose that the heightened OXPHOS in monocytes might reflect increased oxidation of fatty acids and the efflux of cholesterol. Moreover, upregulated genes in neutrophils were predominantly enriched in ribosome-related processes and OXPHOS (Figure 4D and F). The enhanced OXPHOS likely reflected an attenuation of glycolysis-driven inflammation in neutrophils^22^. Consistently, the expression of inflammatory genes (*Il1β, Nlrp3, Ptgs2, S100a8 and S100a9*) in neutrophils showed a downward trend in HFD-26W group (Figure 4E), suggesting a dynamic functional change in neutrophils during atherosclerotic progression.

**Figure 4.**
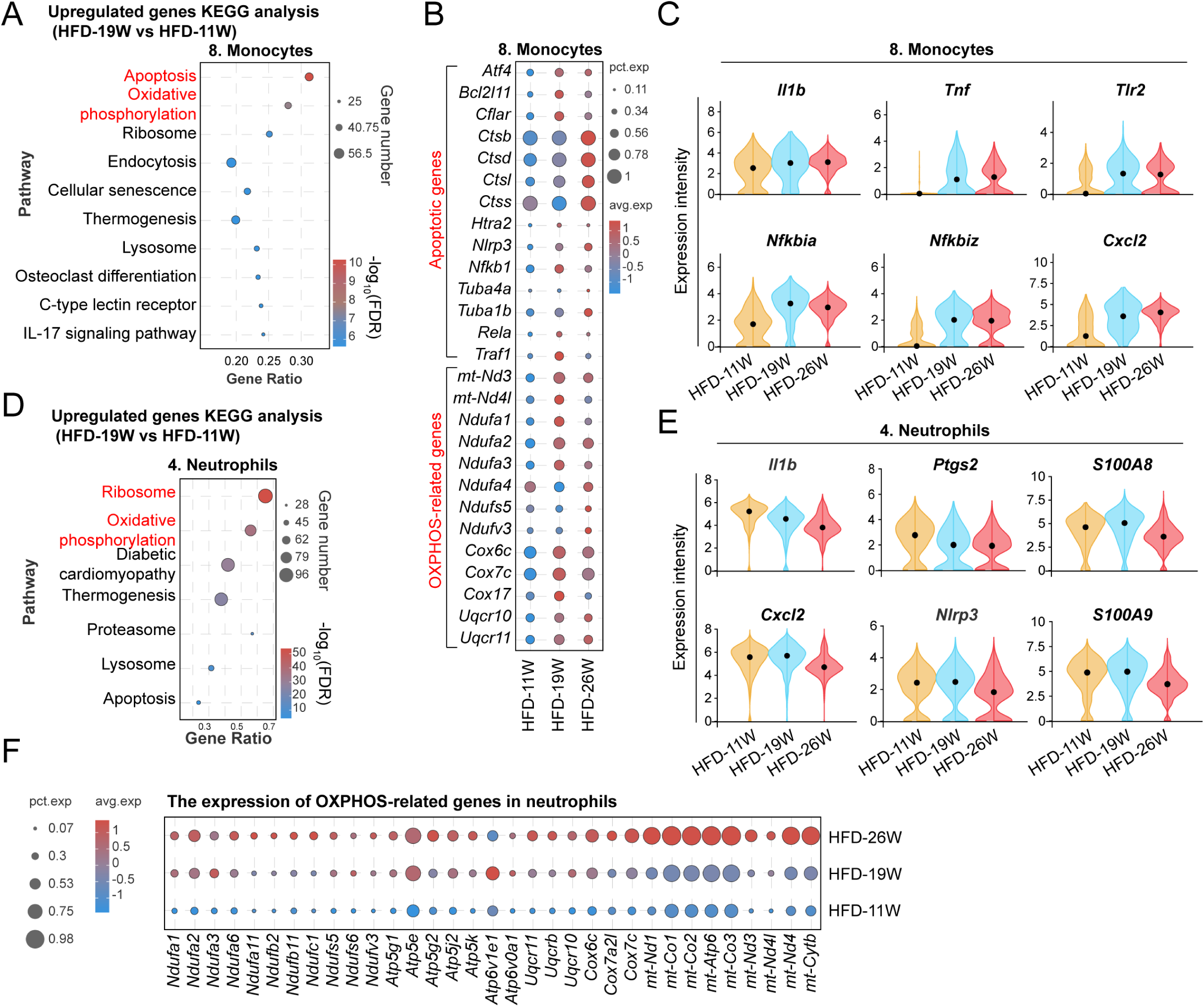
scRNA-seq uncovers functional and metabolic changes in monocytes and neutrophils within the plaque as the disease advances. **(A)** Bubble plot showing upregulated genes enriched signaling pathways in neutrophils (HFD-19 vs HFD-11W) by KEGG analysis. **(B)** Bubble heatmap of normalized gene expression of apoptotic and OXPHOS-related genes in monocytes. **(C)** Violin plot of the expression of specific inflammatory genes in monocytes at different stage of disease. **(D)** Upregulated genes enriched signaling pathways in neutrophils (HFD-19 vs HFD-11W) by KEGG analysis. **(E)** Violin plot of the expression of specific inflammatory genes in neutrophils at different stage of disease. Normalized gene expression. **(F)** Bubble plot showing the expression of OXPHOS-related genes in neutrophils at various stage of atherosclerosis.

### 3.6 Phenotypic transition and functional phenotype of vascular smooth muscle cells during atherosclerosis

Recent studies^4,23-25^ have uncovered the phenotype switch of smooth muscle cells (SMCs) during atherosclerosis. In line with these findings, our analysis of mass cytometric images (Figure 5A), revealed that Collagen Ⅰ in the media region became more abundant as the disease progressed, at the expense of αSMA expression, indicating a phenotypic shift from contractile to secretory phenotype. Our scRNA-seq results supported this notion. As SMC transited to SEM, *Myh11* and *Acta* became downregulated while genes encoding chemokines (*Ccl19* and *Cxcl12*), complement cascades (*C1s1*, *C3*, *C7* and *C1ra*), collagen (*Col3a1*, *Col1a2*, *Col14a1* and *Col1a1*), and mesenchymal stem cell markers (*Gas1*, *Pdgfra*, *Cd34* and *Ly6a*), were greatly upregulated (Figure 5B). Additionally, the differentially expressed genes in the SEM were enriched in the pathway related to protein processing in endoplasmic reticulum and complement cascades (Figure 5C). Through pseudo-time analysis, we identified six distinct differentiation states (SMC1-5 and SEM) from SMC to SME, with SMC1 serving as the initial differentiation point and SEM as “the terminal differentiation state” (Figure 5D). We observed a downward trend in the expression of *Acta2* and an upward trend in *C3*, *Col1a1* and *Col1a2* during the transition from SMC to SEM (Figure 5E). Thus, complement C3 and collagen expression may serve as key indicators of SMC to SEM transition.

**Figure 5.**
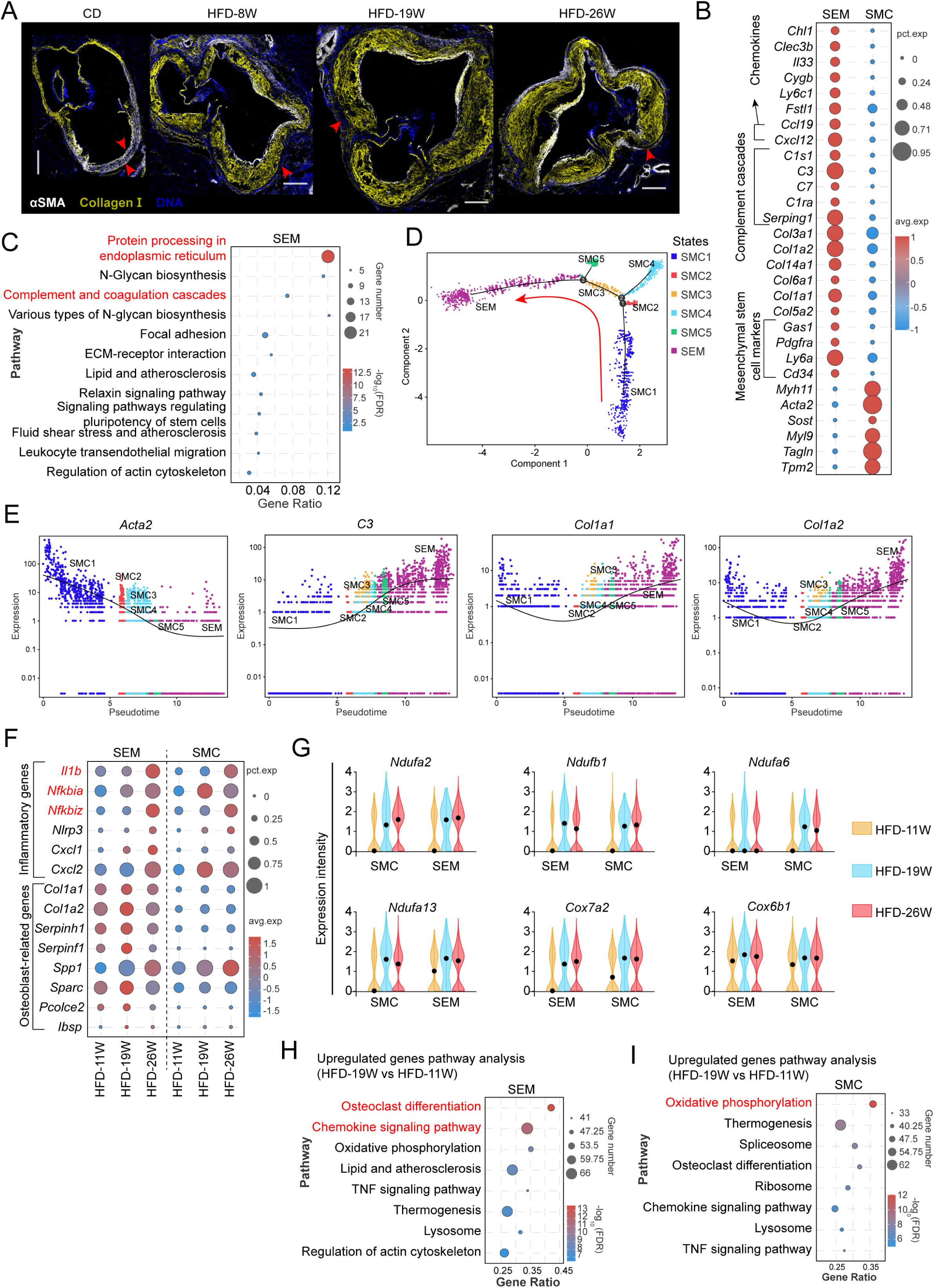
scRNA-seq combined with IMC uncovers phenotype switch and functional changes in smooth muscle cells. **(A)** αSMA and Collagen 1 expression of aortic roots of different groups and red arrows indicating smooth muscle-rich media region. αSMA (white), Collagen 1 (red), DNA (blue), scale bar, 250 μm. **(B)** Bubble heatmap showing differentially genes expression in SEM and SMC populations. **(C)** Differentially genes-enriched signaling pathways in SEM population (Top 12). **(D)** Pseudotime differentiation trajectory plot from SMC transitioned to SEM. **(E)** Pseudotime axis differential genes from SMC transitioned to SEM. **(F)** Bubble plot of the expression of pro-inflammatory and metabolic genes in SEM and SMC population during atherosclerosis. **(G-H)** Upregulated genes-enriched signaling pathways in SEM and SMC (HFD-19 vs HFD-11W) by KEGG analysis. **(I)** Violin plot of mitochondria oxidative phosphorylation-related genes expression in SEM and SMC during atherosclerosis.

Notably, during atherosclerotic progression, both SME and SMC populations displayed higher levels of pro-inflammatory genes, including *IL1β*, *Nfkbiz,* and *Nfkbia* (Figure 5F), as well as OXPHOS-related genes such as *Ndufa2*, *Ndufa6*, *Ndufa13*, *Ndufb1*, *Cox6b1*, and *Cox7a2* (Figure 5G). KEGG analysis of the upregulated genes in SEM and SMC populations (HFD-19 vs HFD-11W) revealed enrichment of pathways related to osteoclast differentiation, OXPHOS and chemokine signaling (Figure 5H and I), suggesting altered energy metabolism and functional changes. Overall, these findings highlight the complex interplay between different subpopulations and the dynamic metabolic rewiring as their functions evolve during the progression of atherosclerosis.

### 3.7 Cell-cell interactions within the plaque microenvironment

To gain a deeper understanding of the spatial plaque microenvironment, we conducted a cell neighborhood analysis using HistoCAT^17^. This analytical approach allows for an unbiased and systematic investigation of all cell-cell interactions present within the plaque and is based on a permutation test to compare the number of interactions between all cell types in the given four images of the aortic root obtained at various stages of atherosclerosis (Representative images derived from 1 x Chow diet, 1x HFD-8W, 1x HFD-19W, and 1x HFD-26W). These distinct pairwise interactions suggest a spatially coordinated plaque microenvironment.

We assigned several cell pairs with significant neighborhood interactions or avoidance (Figure 6A). These pairwise interactions include Adventitia-Mon (cluster 1, 2) interacting with Adventitia-Fibroblast (cluster 5), SMCs (cluster 11 and 15), and T cells (cluster 21); αSMA^hi^ Mac (cluster 8) interacting with Mac nearby fibrous cap (cluster 6 and 16, regarded as foamy macrophages) (Figure 6B) and Neutrophil (cluster 12 and 20); Intraplaque-Mac (cluster 7) interacting with intraplaque-Mon (cluster 23) and αSMA^hi^ Mac; SEM (cluster 12) interacting with Intraplaque-Mac (cluster 7), αSMA^hi^ Mac (cluster 8), and foamy macrophages (cluster 6); Neutrophil (cluster 12) interacting with Intraplaque-Mon (cluster 17 and 23). Notably, many of these interaction pairs were discordant with the pattern of cell frequencies, and exhibited dynamic changes with the progression of disease (Figure 6B-C). Especially, neutrophils-SMCs interactions (cluster12_11, cluster 15_12), were particularly evident during the early-stage and progression of atherosclerotic plaque formation (Figure 6C), which suggested that the interaction between SMCs and neutrophils may be an important element in the formation of early-stage plaque. Additionally, we gained the intriguing observation that neutrophils were predominantly located in close proximity to the smooth muscle cell-rich media region (Figure 2E, Chow diet and HFD-8W). This suggests the potential involvement of SMC in recruiting neutrophils. In combination with scRNA-seq data of aortas, we also predicted cell-to-cell interactions within the plaque macroenvironment through an established computational approach^10^, thereby further supporting these pairwise interactions analyzed by HistoCAT. We observed several significant cell interactions, including SMC-derived SEM-Neutrophils, Gpnmb^hi^-Mac and SMC population, Gpnmb^hi^-Mac-Neutrophils (Figure 6D and E), Adven-Mac-SEM (Figure 6F and G), as well as Inflam-Mac-Neutrophils (Figure 6H and I). These findings support the notion that immune cells and non-immune cells within the plaque intricately regulate plaque formation and state through a complex cellular communication network. For example, neutrophils interact with SEM, SMC, and Mac2 through ligand-receptor pairs such as Fpr1-Anxa1 and Ccr1-Ccl6, potentially contributing to persistent immune response and plaque instability (Figure 6J).

**Figure 6.**
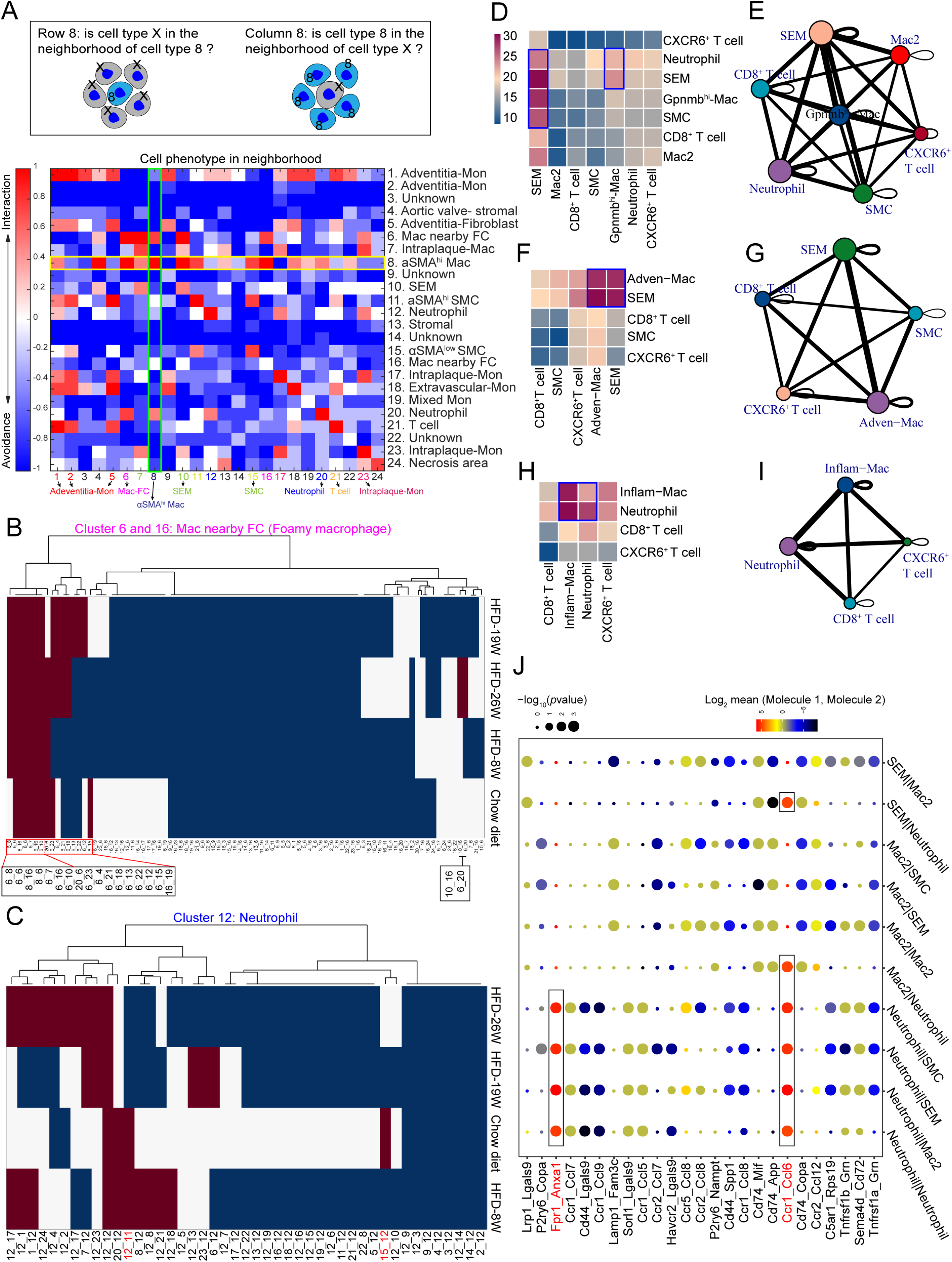
IMC and scRNA-seq reveal cellular interactions within atherosclerotic plaque. **(A) Top,** schematic depicting neighbor interactions visualized in the heatmap. Rows show the significance of all cell types surrounding a cell type of interest. Columns indicate the significance of the cell type of interest surrounding other cell types. White represents an interaction prevalence of less than 10%. **Bottom,** all interactions obtained from 4 aortic root images analysis (Chow diet, n = 1 image; HFD-8W, n = 1 image; HFD-19W, n = 1 image; HFD-26W, n=1 image) are represented as a heatmap in which the cell type in the row is significantly neighbored (red) or avoided (blue) by the cell type in the column. Significance was determined by permutation test (*P* < 0.05). Highlighted squares indicate an example of a directional interaction. **(B-C)** Heatmap showing the significant interaction or avoidance of neutrophil populations between other cell types during atherosclerosis. Significance was determined by permutation test (*P* < 0.05). **(D, F and H)** Heatmap representing the number of predicted ligand-receptor pairs between different cell types within the plaque. **(E, G and I)** Cellular interaction network diagram. The size of each bubble is determined by the number of significantly enriched receptor-ligand pairs between the subtype and all its interacting subtypes. Lines in the diagram represent the number of significantly enriched receptor-ligand pairs between subtypes. **(J)** Bubble heatmap showing the abundance of top 25 ligand-receptor in cell interaction pairs. Dot size indicated the statistical significances. Dot color indicated the average expression of ligand and receptor.

Further analysis of ligand activity by NicheNet^11^ reveals a significant enrichment of the TNF signaling pathway in different cell interaction pairs, including SEM-Adventitia-Mac, SEM-Neutrophil, and Neutrophil-Infam-Mac (Supplemental Figure S4A-C). These observations suggest that the TNF signaling pathway plays a crucial role in cell communication and promotes plaque inflammation.

### 3.8. Single-cell mass cytometry of blood leukocytes reveals increased myelopoiesis during atherosclerosis

The formation of plaques is known to alter hematopoiesis^26^. We utilized mass cytometry to analyze the dynamic immune landscape of peripheral blood during atherosclerosis. Using *t*SNE, we visualized 22 cell clusters, including 8 major immune cell populations (monocytes, macrophages, CD8^+^ T cells, CD4^+^ T cells, B cells, natural killer cells and dendritic cells) based on the expression pattern of canonical markers (Figure 7A-B and Supplemental Figure S5). Of note, during the progression of atherosclerosis, specific myeloid subpopulations (neutrophils in cluster 1 and 10 as well as monocytes in lusters 5, 6, 7, and 12) were significantly increased (Figure 7C). Conversely, there was a notable decrease in B cells (clusters 3, 4, 9, and 11) (Figure 7D), NK cells (cluster 8), and T cells (CD4^+^ T cells, cluster 18; CD8^+^ T cells, cluster 13) (Figure 7E-F). Further analysis showed an elevated neutrophil-to-lymphocyte ratio (NLR) during atherosclerosis (Figure 7G), which is known to be positively linked to cardiovascular disease-related adverse outcomes, including heart failure and ischemic stroke^2,27,28^. Our findings further reinforce the diagnostic or prognostic value of NLR in assessing disease status and the effectiveness of candidate cardiovascular drugs.

**Figure 7.**
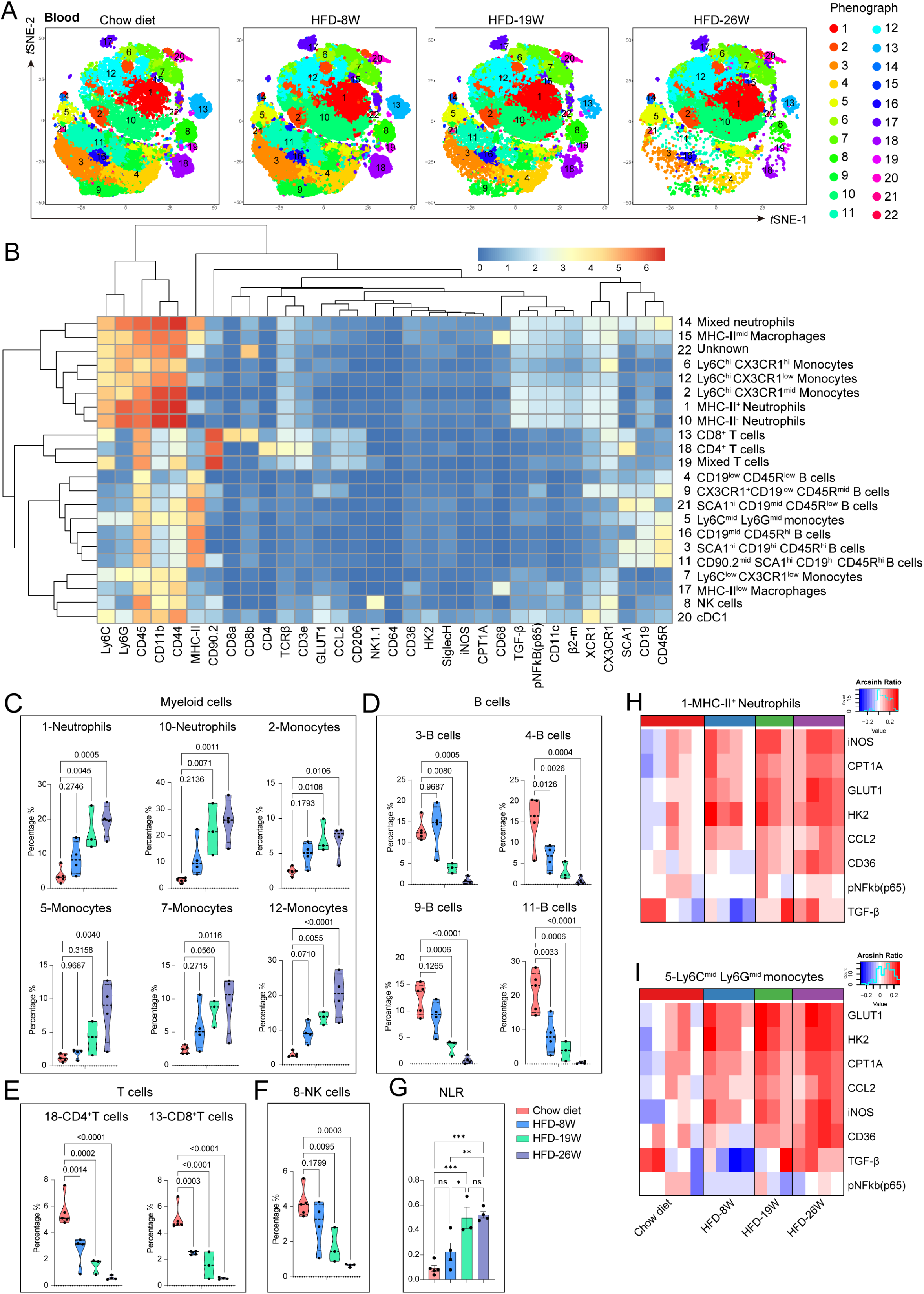
CyTOF reveals the dynamic changes in the circulating immune landscape during atherosclerosis. **(A)** *t*SNE plot of peripheral blood leukocytes from Apoe^-/-^ mice at different stages of atherosclerosis shown the assignment of these cells into 22 clusters. **(B)** Heatmap showing the relative expression level of 32 metal-labelled antibodies within the 22 cell subsets. **(C-F)** Violin plot showing the dynamic changes in abundance of cell populations during atherosclerosis. Myeloid cells (C), B cells (D), Natural killer (NK) cells (E), T cells (F), *p* value was determined by one-way ANOVA with Dunnett’s multiple comparisons test. **(G)** Bar graph showing the dynamic change of neutrophil-lymphocytes ratio at various stages of atherosclerosis. Data are presented as mean ± SEM. Dots represent individual samples, n = 3-5, \**p*<0.05, \*\**p*<0.01, \*\*\**p*<0.001, by one-way ANOVA with Tukey’s multiple comparison test. **(H-I)** Heatmap showing the expression level of glycolysis, lipid metabolism, and inflammation related proteins in cluster 1 and 5 at different stages of disease. The expression levels of antibodies were subjected to Arcsinh transformation, and chow diet was used as a control for comparison.<colcnt=3>

Since physiological functions in organisms are orchestrated by cellular metabolism, we analyzed the dynamic expression of metabolism- and inflammation-related genes in peripheral blood immune cells during the disease process. We selected indicators of glycolysis (Glut1 and HK2), lipid metabolism (CD36 and CPT1A) and inflammation status (iNOS, p-p65 and CCL2). The results revealed that Glut1, HK2, CD36, CPT1A, iNOS, and CCL2 were significantly upregulated in specific cell subpopulations (clusters 1, 5, 10, 11, and 13) in mice on high-fat diet (Figure 7H-I, and Supplemental Figure S6D and I), suggesting hyperactivated glycolysis and lipid metabolism and enhanced inflammatory response in these cell subsets. Additionally, Ly6C^hi^ CX3CR1^mid^ monocytes (cluster 6), defined as inflammatory monocytes, showed upregulation of Glut1, HK2 and p-p65 during atherosclerosis (Supplemental Figure S6B). However, this phenomenon was not observed in other monocyte subpopulations such as clusters 2 and 7 (Supplemental Figure S6A and C), indicating that metabolic and functional rewiring is specific to certain circulating monocyte subpopulations. Despite the significant decrease in CD4^+^ T cells during the progression of the disease, their expression of TGF-β and Glut1 showed a robustly upward trend (Supplemental Figure S6J). This finding could indicate an enhanced immunosuppressive activity in CD4^+^ T cells in response to HFD-induced systematic hyperinflammatory state.

## 4 Discussion

In this study, we present a molecular and spatiotemporal characterization of immune and non-immune cell populations, as well as cell-cell interactions within atherosclerotic plaques using IMC. This approach not only addresses the interference caused by tissue autofluorescence in tissue immunostaining experiments but also mitigates the spatial localization information loss associated with techniques such as scRNA-seq and CyTOF, which depend on tissue enzymatic digestion. Moreover, many plaque cell populations identified using IMC correspond to aortic cell subsets identified through scRNA-seq analysis of aorta, such as Adven-Mac, Gpnmb^hi^-Mac, Inflam-Mac, SMC, SEM, and neutrophils. Thus, IMC can complement other technological approaches in delineating the dynamic changes in the composition and distribution of plaque cells during atherosclerosis.

In this study, we observed elevated frequencies of Intraplaque-macrophages, Intraplaque-monocytes, Adventita-Mon and SEMs in atherosclerotic plaques, which was accompanied by a reduction in contractile SMCs. These findings were consistent with previous studies^4,7,29^. Notably, our results indicate that αSMA^hi^ Macs, located in the fibrous cap of atherosclerotic lesions, do not expand with disease progression and may predate early-stage plaque formation, suggesting their independent role in determining the fibrous cap thickness. IMC also allowed us to locate a significant number of neutrophils in aortic root, which peaked at the HFD-8W (intermediate-stage), indicating that neutrophil infiltration may contribute to atherogenesis and inflammation. scRNA-seq analysis revealed that these neutrophils display a hyperinflammation phenotype, but the expression of inflammatory genes did not further increase with disease progression. Cell neighborhood analysis by IMC uncovered that neutrophils in early-stage plaques primarily interact with SMC and αSMA^hi^ Mac populations, but with Intraplaque-Mon and Mac in late-stage plaques. This suggests that the neutrophils may have been recruited by activated smooth muscle cells and macrophages in early-stage plaques, and form neutrophil extracellular traps to sustain continuous immune responses and plaque formation^30^. Using ligand-receptor interaction analysis, we further confirmed the interaction between neutrophils and SMC, SEM, and foamy macrophage likely via CCR1-CCL6^31^ and FPR1-ANXA1^32^.

Cellular metabolic states of plaque cells are intricately linked to their functions, thereby influencing atherosclerotic progression^33-35^. Here, we found that glycolysis, OXPHOS and electron transport chain were all more active in Gpnmb^hi^-Mac than in Adven-Mac and Inflam-Mac, in contrast to the conventional glycolysis-M1 and OXPHOS-M2 associations. Next, we characterized the alterations of plaque cells in biological functions and metabolism-related signaling pathways from early-stage plaques to advanced plaques using KEGG analysis. We observed activated chemokine and TNF signaling pathways, as well as enhanced OXPHOS, in macrophage populations, SMCs, and SEMs. Although OXPHOS is generally considered as an anti-inflammatory metabolic characteristic in monocytes/macrophages, in the context of atherosclerosis, the enhanced OXPHOS in macrophages may be an adaptive strategy in response to excessive lipids that need to be handled by fatty acid oxidation to reduce lipid accumulation and ER stress in macrophages. However, boosting OXPHOS and osteoclast differentiation in SMCs and SEM was specifically associated with intensified vascular calcification^36^.

The pathological changes caused by atherosclerosis are not limited to local vascular plaques but also generate systemic inflammatory response^37-39^. Through high-dimensional CyTOF analysis, we delineated the dynamic immune landscape in peripheral blood, and observed elevated myeloid cells and decreased lymphocytes, accompanied by activation of metabolic and inflammatory pathways. Inhibiting TNF signaling pathway with special neutralizing antibody may potentially hinder disease progression and reduce the occurrence of cardiovascular events, which is supported by two clinical trials^2,40^ showing therapeutic effect of IL-6 inhibition with ziltivekimab and IL-1β inhibition with canakinumab, respectively.

Our study has several limitations. Firstly, we were unable to identify dendritic cells, CD4^+^ T cells, CD8^+^ T cells, and vascular endothelial cells in the plaque due to lack of suitable metal-conjugated antibodies. Our attempts to classify macrophages into anti-inflammatory or pro-inflammatory subtypes were also limited by the low expressions of Arg1 and iNOS. Secondly, our study focused on experimental mice with atherosclerosis, and there are inherent species differences between mice and humans in the cellular composition of plaques. For instance, human plaques are characterized by abundant T cells, whereas mouse plaques are predominantly composed of macrophages. Therefore, the findings from our study may not fully reflect the characteristics of human plaques. Thirdly, we did not validate the observed metabolic and functional changes in the identified cell subsets at the protein level. Further examination at the protein level would enhance our understanding of the specific metabolic and functional changes in these cell subpopulations within both human and mouse atherosclerotic plaques.

In summary, we generated a comprehensive atlas of murine atherosclerotic plaque, including the spatiotemporal dynamics, cell interaction networks, molecular features. The new information may improve our understanding of the underlying mechanisms of atherosclerosis and help identify novel therapeutic targets for the treatment of cardiovascular disease.

## Author Contributions

P.H., J.F., and Z.L. conception and design, collection and assembly of data, animal experiments, data analysis and interpretation, and manuscript writing; S.W., Z.W and S.L. contributed to analytic tools, antibody conjugation and IMC experiments; G.M provided constructive feedback on the paper’s writing; P.L., C.S. and Y.S.: conception and design, manuscript writing, administrative support and financial support.

